# Diversity of Archaea Domain in Cuatro Cienegas Basin: Archaean Domes

**DOI:** 10.1101/766709

**Authors:** Nahui Olin Medina-Chávez, Mariette Viladomat-Jasso, Gabriela Olmedo-Álvarez, Luis E Eguiarte, Valeria Souza, Susana De la Torre-Zavala

**Affiliations:** Universidad Autónoma de Nuevo León, Facultad de Ciencias Biológicas, Instituto de Biotecnología. Av. Pedro de Alba S/N Ciudad Universitaria. San Nicolás de los Garza, Nuevo León, México. C.P. 66455; Instituto de Ecología, UNAM, Circuito Exterior S/N anexo Jardín Botánico exterior. Ciudad Universitaria, Ciudad de México, C.P. 04500; Departamento de Ingeniería Genética, Centro de Investigación y de Estudios Avanzados del I.P.N. Campus Guanajuato, AP 629 Irapuato, Guanajuato 36500, México

## Abstract

Herein we describe the Archaea diversity in a shallow pond in the Cuatro Ciénegas Basin (CCB), Northeast Mexico, with fluctuating hypersaline conditions containing elastic microbial mats that can form small domes where their anoxic inside reminds us of the characteristics of the Archaean Eon, rich in methane and sulfur gases; thus, we named this site the Archaean Domes (AD). These domes only form after heavy rains that are rare in the Chihuahuan desert. CCB is a unique oasis with hundreds of ponds, containing endemic species of animals, plants and highly diverse and unique microbial communities, despite its very biased stoichiometry, due mostly to extreme low phosphorus content (soils, water columns and sediments). This extreme oligotrophy has favored survival of ancestral microorganisms. Whole metagenome sequencing approach was performed for this unusual site in three different seasons to assess the extent of the Archaea biodiversity, with a focus on extremophiles, since members of the Archaea had been underrepresented in different study sites within the oasis. We found a highly diverse Archaea community compassing ∼5% of the metagenomes. The archaeal portion in all three metagenomes maintained its abundance and most of the strains showed to form a resilient core during three seasonal samplings (2016-2017), despite environmental fluctuations. However, relative abundances of all 230 archaeal OTUs (defined using a 97% cutoff) were low enough (<0.1%) to be considered part of the rare biosphere. AD finding and their description within CCB confirms that this particular pond is the most diverse for Archaea that we are aware of and opens new paths for understanding the forces that once drove and keep shaping microbial community assemblage.

## Introduction

The Archaea domain is an essential but usually rare, component of different ecosystems, not only extreme ones as originally suggested [1–4], for instance, they are critical in hydrological systems and their abundance and composition changes according to identifiable spatial and temporal scales [5–7]. Archaea is represented in 26 phyla [8], most of which are novel and some lineages include only one species of uncultivated strains that have been assembled through metagenomics [9, 10]. Indeed, a current challenge in their study is the isolation and cultivation of novel Archaea, since they are usually restricted to extreme conditions. Traditional culture-dependent approaches have underestimated abundance and diversity of Archaea, making it hard to achieve an accurate understanding of their role in microbial communities in an ecological niche [11]. Nevertheless, a successful co-cultivation of an Asgard archaeon associated with bacteria was reported after a long-term methane-fed bioreactor culture of deep marine sediments, demonstrating that non-traditional cultures of untapped environments are reservoirs of Archaea genetic and functional diversity yet to be uncovered [12]. Another challenge is to fully comprehend their phylogenetic association, given the limited genetic information available for some Archaea phyla.

In the last decade, our knowledge on the diversity and taxonomy of Archaea has substantially improved [13] due to the rise of metagenomics and increasing available archaeal genomes. The Archaea tree is being rapidly filled up with new branches, demonstrating that the Archaea domain remains largely unexplored [14] along with a diverse metabolism [15, 16], unveiling new processes and key features involving microbes and community structure.

Cuatro Cienegas Basin (CCB) is an endangered oasis within the northern zone of the Chihuahuan Desert in Mexico, characterized by an extremely unbalanced nutrient stoichiometry of the area (N:P = 159:1), similar to the conditions of the Precambrian sea [17–21]. Strikingly, despite this nutrient deficiency CCB is considered a biodiversity hotspot for macroorganisms [22] and one of the most diverse sites for microorganisms in the world [23–27]; this microbial biodiversity is mirrored by its extreme diversity in virus [28]. CCB biodiversity is believed to have evolved as a result of a long time environmental stability of a deep aquifer [29], as suggested by the marine affiliations in many of the studied bacterial genomes [25, 30], virus [31], and probably, Archaea. These observations led CCB scientists to raise hypothesis and propose CCB as a model of “*lost world*”, where extreme conditions favored the survival of ancestral marine lineages that in some cases persist exclusively in this area [32]. Nevertheless, Archaea in CCB has been poorly studied since they were underrepresented in previous metagenomic studies [24, 28, 33, 34].

During a late March 2016 field work, an atypical spring rain apparently dissolved the salty crust of an unnoticed shallow pond, allowing the uniquely flexible and impermeable microbial mats to arise from the ground building bubbles, or domes with a strong sulfur-like smell. Our hypothesis was that gas production of methanogenic as well as other microorganisms associated to the sulfur cycle were degassing creating a locally anaerobic atmosphere. Therefore, the anaerobic layer of the microbial mats were creating the domes by raising the flexible and impermeable mat where photosynthesis was evident given the purple, and green layers, creating an unique “alien” local landscape (Figure 1). Based on the macroscopic morphology of the microbial mats, this locality in the Pozas Azules ProNatura ranch (a private ecological reserve) was named by us the “Archean Domes” (AD thereafter) (Fig. 1a).

**Figure 1.**
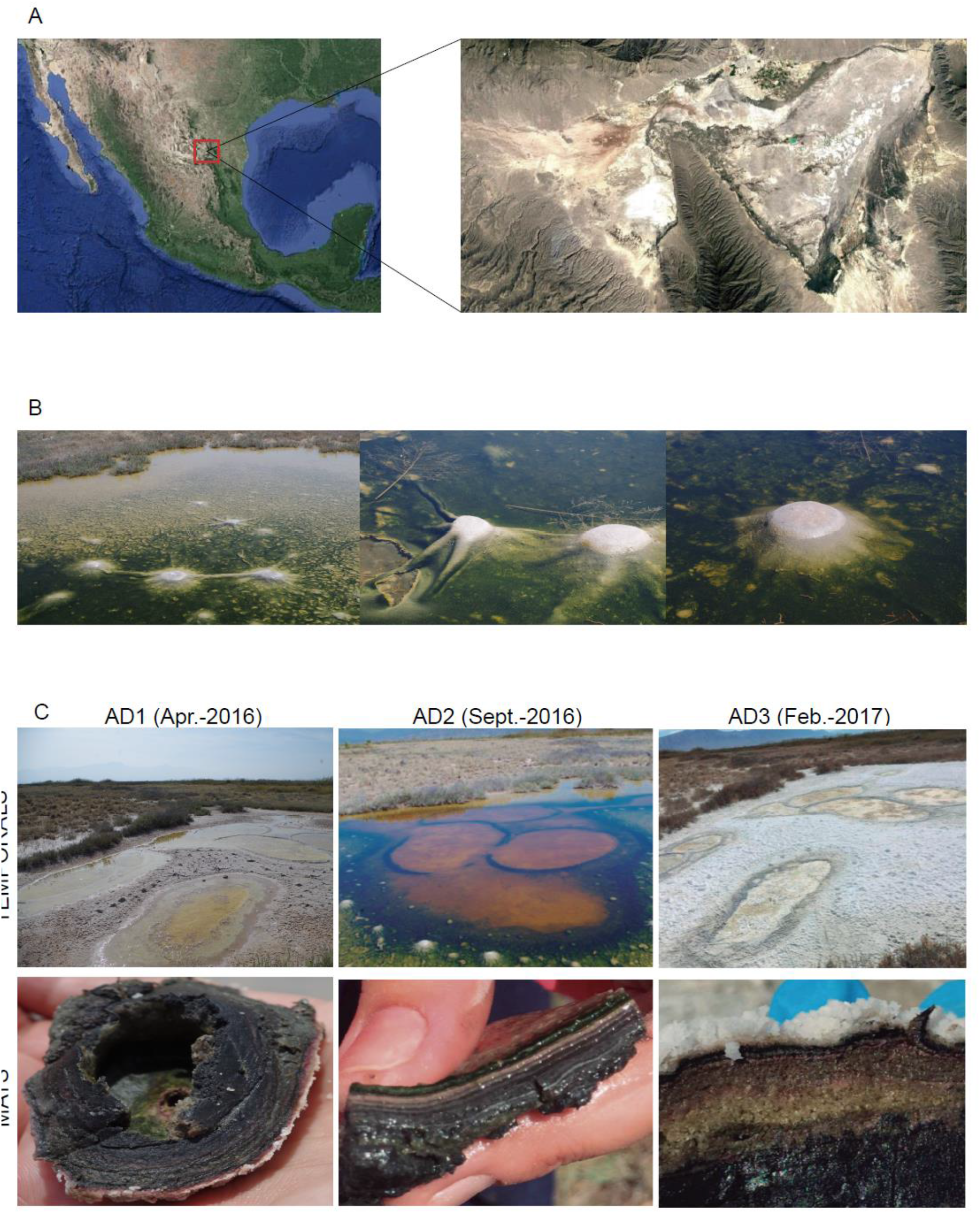
Sampling site Map. A) Cuatro Cienegas, Coahuila location within Mexico; B) Archaean Domes (AD) mats with unusual and flexible bubble-containing superficial layer found in Pozas Azules Ranch within Chihuahuan Desert region C) AD sampling of the three continuous seasons: AD1 (April 2016), AD2 (September 2016), AD3 (February 2017) showing their obtained microbial mat and the structure of microbial community in in its respective season.

These unusual structures in an small extremophilic and fluctuating pond, are by themselves a singularity, within CCB unique nature and amazing microbial diversity that has fascinated scientists with unexpected findings [19-21, 25, 27, 29, 30, 33, 35-43]. High salt concentration in dry conditions, and the production of biogases along dome formation under wet conditions, suggested that this fluctuating evaporitic hypersaline microbial mat would be an ideal place to explore the Archaea diversity in CCB.

Considering the taxonomic profiling of different ponds previously studied within the Cuatro Cienegas system along 20 years [17, 28, 30, 33, 37–39, 41, 44–53], a working hypothesis was proposed for the community composition of AD microbial mat: that each sample would be unique, given the drastic environmental fluctuations. A null hypothesis would predict that the AD community would constitute a unique community with a constant core community, able to construct the domes given the right environmental conditions (heavy rain). In order to explore these hypotheses for the Archaea domain, a whole genome metagenomic approach was applied to explore the community shaping the AD microbial mat, expecting to achieve a coverage depth sensitive enough for the rare biosphere. Sampling for AD was done in three seasons (dry-wet-dry).

As mentioned above, Archaea had not been observed to be diverse or abundant in CCB, however, we found in this study that the Archaean Domes displayed an abundant and diverse archaeal community. Even more, they represent one of the most diverse archaeal communities known up to date in the world. Unexpectedly, this diversity is not only very large given the small geographical scale of this site (at cm^2^ level), but it is constant along all sampling seasons. This novel window to the Archaea world will be a biodiversity resource that may provide us with paths to understand the uniqueness that CCB *lost world* represents for ecology, evolution, systematics and future bioprospecting studies.

## 2. Material and Methods

### 2.1 Sampling site

Water and sediment were obtained from the Archaean Domes (AD) in Rancho Pozas Azules, from Pronatura Noreste (26° 49’ 41.7’’ N 102° 01’ 28.7’’ W) in CCB, under SEMARNAT scientific permit No. SGPA/DGVS/03121/15 (Figure 1). Sampling took place during 2016-2017 at the following times: April 2016 (dry, AD1), September 2016 (wet, AD2), February 2017 (dry, AD3); microbial mats and sediments (8 cm^2^) were transferred to sterile conical tubes (50 mL) and used for metagenomic analysis.

### 2.2 Metadata

Following the National Meteorological System data base for the Pozas Azules ranch weather station (Supplementary material), we observed that in March 2016, there was an anomalous spring rain peak (90 mm) that apparently allowed us to discover the AD. The 10^th^ of April sampling occurred when maximum air temperatures where ∼ 30° C, and the domes were drying. In average, precipitation in early fall, reached 48 mm in September 2016, during our wet sampling date, and no rain were detected during February 2017, making it another dry sampling. The water physicochemical conditions were measured in September 2016 sampling, using a Hydrolab MS5 Water multiparameter sonde (OTT Hydromet GmbH, Germany) (Supp. Table 1). Other sampling dates were not tested due to the lack of water in the ponds (see Figure 1).

### 2.3 Total DNA Extraction

Extraction of total DNA from Archaean domes sediment was performed in Experimental and Molecular Evolution Laboratory of the Ecology Institute, UNAM as reported before [24] using a modification of Purdy et al. (1996) [54] protocol.

### 2.4 Metagenomic Shotgun Sequencing

Total DNA from sediment of the three seasoned samples (AD1, AD2, and AD3) were sent to CINVESTAV-LANGEBIO, Irapuato, Mexico, for shotgun whole genome sequencing using Illumina Mi-Seq 2×300 technology.

### 2.5 Bioinformatic analysis for metagenomes

Raw data from AD1, AD2 and AD3, was quality checked using FastQC (http://www.bioinformatics.babraham.ac.uk/projects/fastqc). Indexed adapter and barcodes were removed and low quality sequences were discarded with Trimmomatic v0.36 using a sliding window of 4pb and an average quality per base of 25 [55], followed by the merging of the paired end reads using PEAR software [56]. Once reads were merged, both paired ends assembled and unassembled reads were used to cover major accuracy to obtain relative abundance for Eukarya, Bacteria and Archaean species using MetaPhlAn2 software [57].

### 2.6 Diversity analysis and species accumulation curve

Using normalized relative abundances of Archaean taxa within each metagenome, α and β diversity were calculated using EstimateS 9.1.0 and Past Software, respectively [58, 59].

### 2.7 Phylogenetic analysis

Reference 16S rRNA sequences were downloaded from the NCBI-*Gen Bank* and *RNA Central* Database according to identifications of MetaPhlAn2 hits. The sequences were aligned with ClustalW [60] and trimmed using MEGA V 7.0 [61]. The phylogeny was reconstructed with a Maximum Likelihood (ML) algorithm in MEGA and K-2 + G (Kimura 2-parameter + gamma distribution) evolutionary model with 10,000 bootstraps. Also reference strains were added to supply a robust interpretation.

### 2.8 Identification number of Metagenomic Data

The identification number for the metagenomic data used in this study are available in MG-RAST with the following ID: AD 1: mgm4856917.3, mgm4856915.3; AD 2: mgm4856913.3, mgm4856914.3; AD 3: mgm4856916.3, mgm4856918.3, and also using the following link: https://www.mg-rast.org/linkin.cgi?project=mgp90438.

## 3. Results

### 3.1. The “Archaean Domes” are hypersaline non-lithifying microbial mats

After obtaining the necessary collecting permits and adequate equipment, sample collection of the AD was conducted in early April 2016. By then, the AD site was starting to dry, and the mat exhibited a morphology in which a photosynthetic upper layer became evident in the dome-like structure, while the interior of the domes maintained a wet black layer (Fig. 1b). Physicochemical and environmental conditions of AD were registered on April 2016, September 2016, and February 2017, that is, during dry and wet conditions (Table 1 and Supp. Table 1). Further observations (not shown) after four years revealed that during the dry seasons, AD microbial mats are active even if they do not produce domes. Moreover, it is only after heavy rains dissolve the salts, that domes emerge due to the elastic nature of the mats, producing its typical sulfur-like smell. We have observed the same phenomena up to last exploration September 2019.

**Table 1.** Physicochemical parameters and Archaeal abundance in four Hypersaline Microbial Mats.

| Microbial Mat | Salinity % | Temperature C° | pH | Archaea relative abundance % | Archaeal OTU's (97%) | Sequencing approach | Reference |
| --- | --- | --- | --- | --- | --- | --- | --- |
| Archaeal Domes | 53% | 31.45 | 9.94 | 4.96% | 230 | Whole Genome-Shot Gun Illumina MiSeq | This study |
| Guerrero Negro, Mexico | 28% | 30 | N/A | 30.45% | 209 | 16s rRNA Tag (926F/1392R) FLX Titanium Roche | García-Maldonado *et al.*, 2018 |
| Shark Bay, Australia | 6.8% | 24.8 | 8.13 | <1% | 13 | Whole Genome-Shot Gun (Illumina MySeq) | Babilonia *et al.* 2018 |
| Atacama, Chile | 10.6-15% | 23-27 | 7.65 | 50-66% | 121 | 16s rRNA Tag (F515/R806) 454 pyrosequencing | Fernandez *et al.* 2016 |

The AD mats sampled on April 10^th^, 2016, looked different from the original observation (less than 3 weeks earlier). What firstly seemed like a shallow pond, was already drying up when the first sample was collected for sequencing (AD1), and the dense salty liquid started to turn into a salty-white-carbonate crust. As can be seen in Figure 1C, in, the second sampling (AD2, September 2016) represented a more wet environment, and the site looked similar to the one observed originally (i.e., March 2016), although the domes were smaller. The last sample of this study was collected in February 2017, by the end of the cold-dry season (AD3).

The microbial community of the AD is approximately 200 m away from the most described, stromatolite-rich and iconic blue pond from Rancho Pozas Azules, but physicochemical conditions found in the AD are extreme in salinity and higher in pH (Table 1). While Blue Pond pH is 7.9 [62, 63], pH of AD almost peaks 10. Moreover, in the first sampling (AD1), salinity was 53%, and it reached saturation later in that year, when the upper layer of the mat became a white salty crust, which prevented measurement with the available equipment for consistent results. Considering all the above, we can describe the AD as a hypersaline microbial mat [64].

### 3.2. The Archaea domain in the Archaean Domes

The Illumina MiSeq run of 300-bp paired-ends delivered 28,859,454 reads for AD1, 24,772,053 reads for AD2 and 28,203,484 reads for AD3 of total DNA metagenomics, which were normalized for bioinformatic analysis. The percentage for relative abundance of the Archaea domain in AD1, AD2, and AD3, were 3.6%, 5% and 5% respectively. The binning of the metagenomic reads showed that the archaeal community in the AD metagenomes was dominated by Euryarchaeota-halophilic archaeal lineages. Sequences from Crenarchaeota, Thaumarchaeota, Korarchaeota, and Nanoarchaeota phyla were also retrieved (Fig. 2). All five phyla comprise 15 classes dominated by members of Halobacteria and Methanomicrobia classes (Fig. 2), both important groups of the Euryarchaeota phylum. Additionally, 25 orders, 36 families, 93 genera, and 230 *species* (defined as OTUs at 97% cutoff) were detected in the three metagenomes. It is noteworthy that from those 230 OTUs, a total of 24 could not be further classified (Supp. Fig.2).

**Figure 2.**
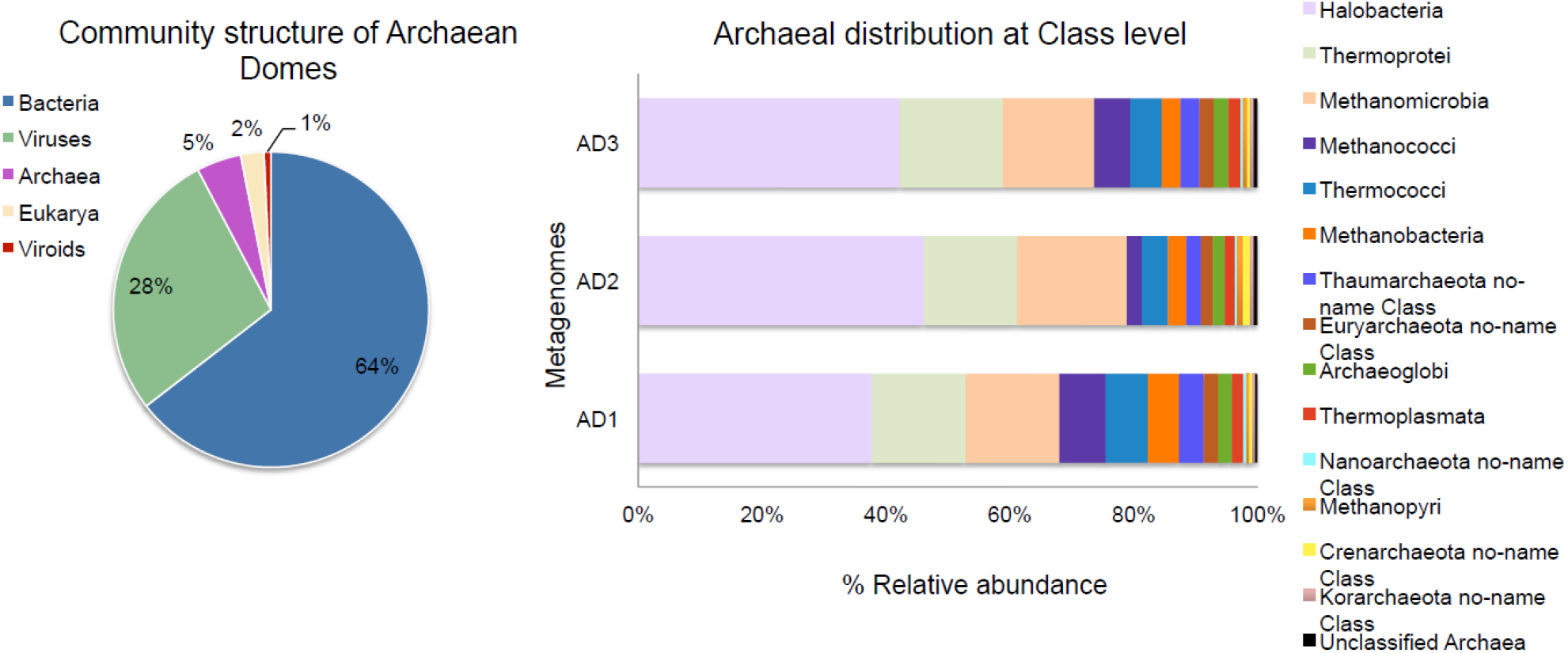
Metagenomic data of Archaean Domes. Mean Community composition (left) and Class-Level distribution (right) of Archaea in the three continuous season samples of Archaean Domes.

To depict the extent of the biodiversity present within the AD metagenomes, we retrieved from the marker databases the 16S rRNA sequences corresponding to the species identified by MetaPhlAn2. We used these sequences to reconstruct a phylogeny (Fig. 3). Several sequences belonging to the same genera were omitted to avoid redundancy; others without a full-length marker sequence were also omitted. The phylogenetic reconstruction displays 213 sequences, 169 belonging to our mining and 44 are reference sequences. Five phyla were represented, that gave rise to 11 major clades.

**Figure 3:**
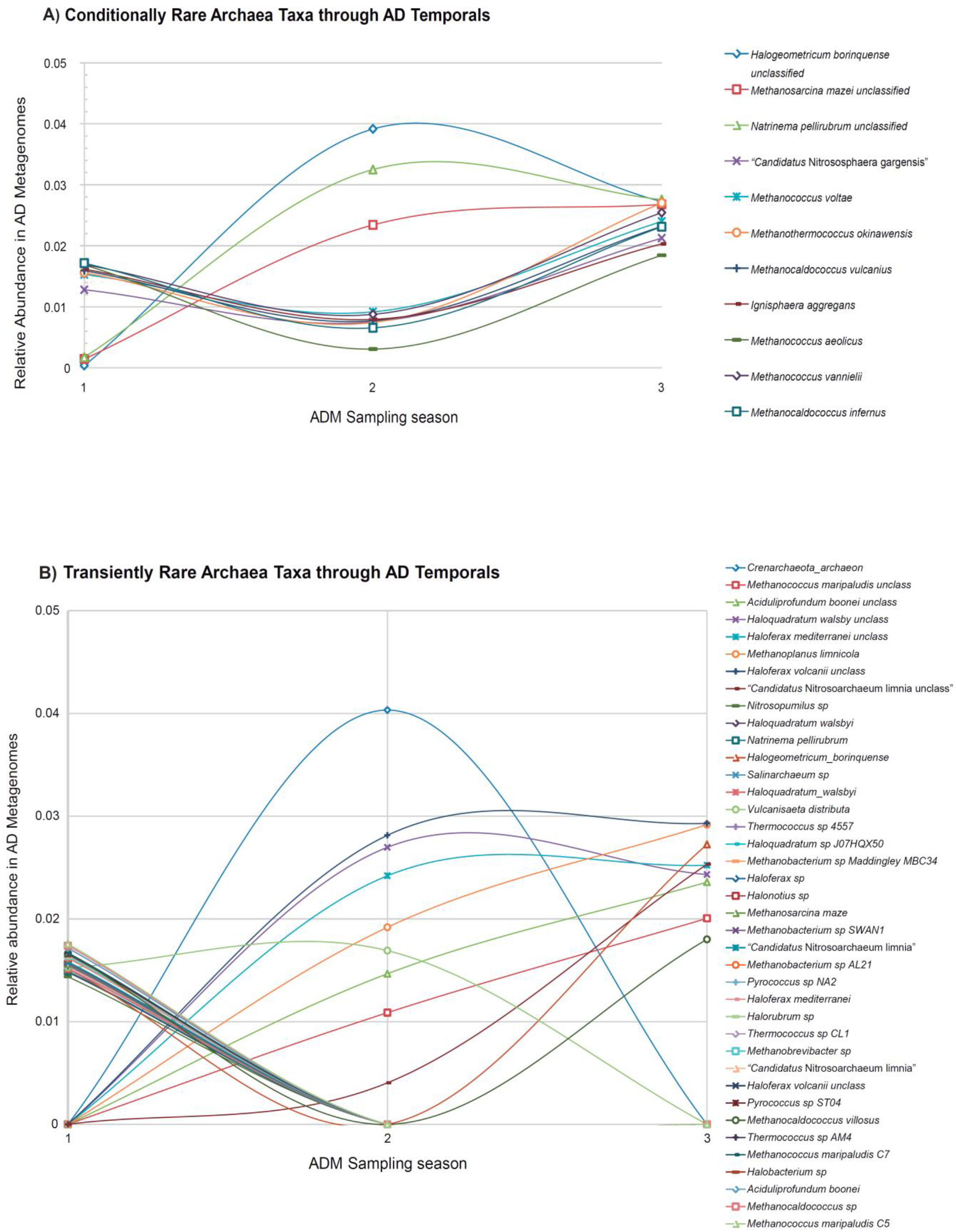
Archaean rare biosphere. A) Relative abundance of Conditionally Rare archaeal taxa through seasons in the Archaean Domes. B) Relative abundance of the Transiently Rare archaeal taxa through seasons in the Archaean Domes.

**Figure 4.**
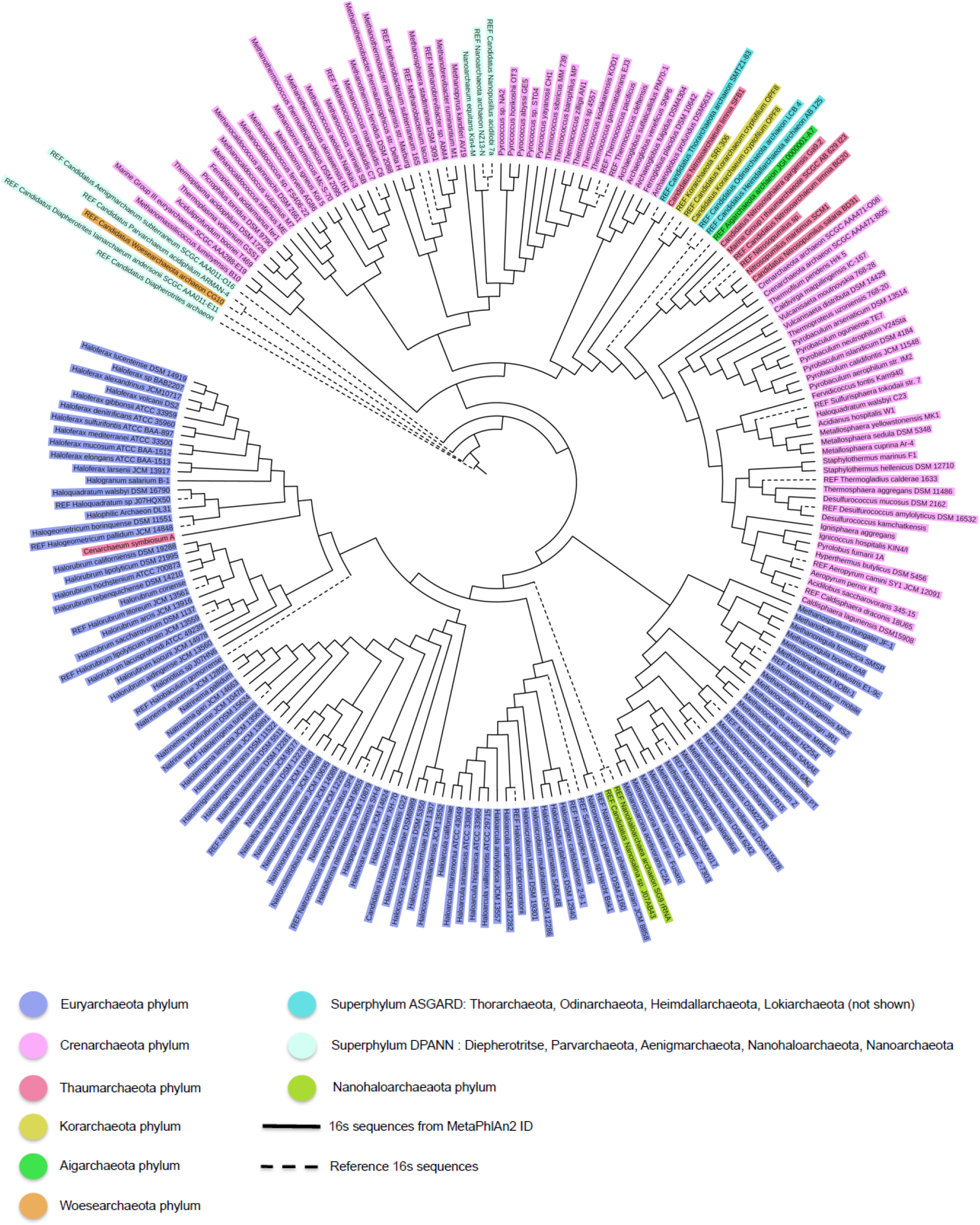
Phylogenetic tree of the Archaea domain species found in AD1, AD2, AD3, though MetaPhlAn2 hit profile, based on nearly full-length 16S rRNA gene sequences, using maximum likelihood method, constructed by K-2 + G evolutionary model with 1,000 bootstraps replicates.

Alpha diversity was calculated using Shannon, Simpson and Chao indexes (Table 2a). Compared to the fluctuations observed in several bacterial studies in CCB [65], the archaeal diversity did not show drastic changes through sampling seasons (Table 2a). Moreover, 90% of the total archaeal richness (OTUs at 97% cutoff level) is shared among the three samples. This is confirmed by the Beta diversity estimates at species level (Table 2 b). Overall, richness evaluated at the other different taxonomic levels (i.e., using the 15 classes, the 25 orders, and/or the 36 families) of Archaea, remain similar in the three metagenomes.

**Table 2.**
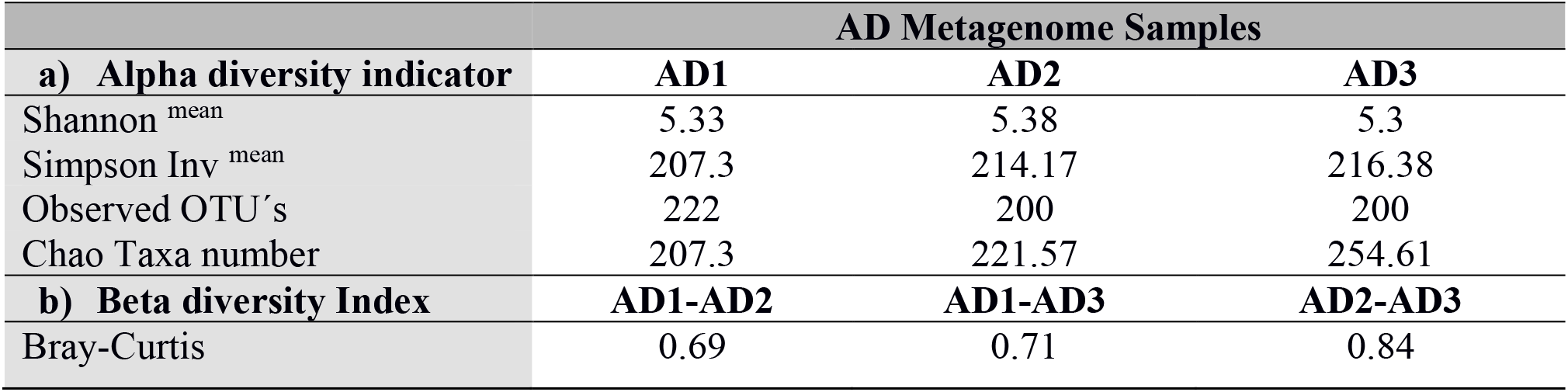
Diversity indexes of the Archaean Dome Metagenomes. a) Alpha and b) beta estimator indexes for specific Archaea diversity.

In accordance with the Archaea’s stable core hypothesis for the AD, the species accumulation curve shows that the inventory for every sample (AD1, AD2, AD3) was very similar. The expected diversity using the presence and absence for every identified OTUs in the metagenomic analysis increased very little with the addition of each sample. Therefore, OTU rarefaction curve with data from the 3 samples is asymptotic (Suppl. Fig. 1).

### 3.3 The Archaean Domes Rare Biosphere

Considering that 229 of the 230 identified archaeal OTUs are each present at relative abundances below 0.1%, we applied a strict threshold of 0.01% of relative abundance [66] to identify what we consider *strictly rare* archaeal taxa. Consequently, within the AD mats, 50 archaeal OTUs were defined to be strictly rare (each covering less than <0.01% of the total reads). Interestingly, such abundance profiles along the three sampling times show that 11 OTUs of the strictly rare Archaea taxa seem to be *conditionally rare* [67], that is, these taxa maintain their lower abundance in only one sample, and reach higher (while still rare <0.1%) abundances in other samples (Fig. 3A). This suggests that while the rest of the Archaea community is relatively stable, the strictly rare biosphere is dynamic and depends on the proper conditions to prosper. On the other hand, we found that 39 strictly rare Archaea OTUs were not detected in at least one metagenome (Fig. 3B). These taxa exhibiting this kind of fluctuation along time series are considered *transiently rare* taxa [68]. Notice that sample season 2 (wet) shows a peak of transient rare taxa, suggesting that it is possible that dome construction during wet conditions can benefit by this change in population dynamics, while other taxa seem to prosper in the extremely salty environment.

## 4. Discussion

The microbial community that constitute the AD microbial mats is indeed unique among the studied CCB bacterial communities [21], displaying a relative high abundance of Archaea in AD1, AD2, and AD3, that reached 3.6%, 5% and 5% respectively. Those abundances are remarkable for such a small sampled area (cm^2^), but also when compared to other sites in CCB and the world (see Table 1) [69–71]. After profiling the community at several taxonomic levels, it was observed that an archaean community core remained almost intact in time (April 2016-February 2017). This provided evidence for stability behind the construction of the domes (our null hypothesis). Nevertheless, within that core, the diversity and proportions of rare archaeal taxa increased or diminished across seasons, exhibiting a dynamic in the AD which might be playing a role for their functioning along different environmental conditions.

### 4.1 The Archaean Domes at CCB are a good model of the Archaean period, and are one of the most abundant and diverse environments for Archaea

Microbial mats and stromatolites are considered analogues of early Earth, when they originated as successful ecological communities, capable of complete nutrient cycling as soon as life started to diversify in the early Archean Eon [72, 73]. Different ancestral lineages have survived at CCB [21, 30], purportedly because they have adapted continuously to their neighbors, forming very tight communities that cohesively survive despite the changing harsh abiotic factors, like those experienced along their very long evolutionary trajectory [21]. We believe, that the microbial mat of AD is not an exception in CCB, however, the observation that raised scientific interest to study these particular mats was precisely its extreme conditions of pH and salinity, besides their interesting “architectural” shape. Notwithstanding, analogue extreme environmental conditions have been reported in other archaeal rich microbial mats before, for instance in the Desert of Atacama or the human made salterns in Guerrero Negro, both stable environments (Table 1) show much lower Archaea diversity [69–71]. A possibility is that the fluctuating environment in AD are part of the community dynamics increasing possible niches, and it has been observed within CCB in other fluctuating environments that harbor large diversity due to local adaptations to each condition [37, 74]. However, the AD mat is peculiar because Archaea are more abundant and diverse than in other fluctuating microbialites within CCB [62] (Table 1). Moreover, this is the first site where there is a constant microbial core despite the environmental fluctuations among seasons. Not only Archaea diversity within AD is relatively constant, it is also considerably large for a metagenomic study using the Illumina platform [75].

On the other hand, targeted clone libraries for Archaean primers have revealed a high abundance of Archaea (∼10%) in stromatolites communities of the Hammelin Pool in SharkBay, Australia, which reaches one of the highest archaeal abundances. However, such archaeal community was represented by 27 archaeal clones, out of 176 total of 16S rRNA clone library from a single sample point [76]. These numbers are hard to compare to our observed AD archaeal diversity due to different experimental approaches. Other archaeal-rich hypersaline microbial mats show relative abundances of Archaea that range from 4% in a mat in Camargue, Spain. [77], 9% in Guerrero Negro hypersaline mat [78], 20% in some layers of the microbial mat in a shallow pond in the Salar de Llamara [79], and above 90% of 16S rRNA of sequences amplified in a highly lithified mat in Laguna de Tebenchique, in the Salar de Atacama [69]. Nevertheless, all these Archaea-rich sites, display lower diversity [80] than our small (cm^2^) seasonal AD. Given the extremely small site, our 230 different species (at 97% cutoff) and 5 different phyla can be considered high. Noticeably, our 24 unclassified archaeal OTUs (Supp. Figure 2), constitute an interesting challenge for systematics and biodiscovery.

The CCB is well known for its outstanding microbial diversity and unique endemic lineages [25, 28, 35, 40, 74, 81] that have been isolated from the rest of the world; this uniqueness may be in part the result of its extremely biased stoichiometry [17, 19, 44]. Success of archaeal communities in the AD may rely on physiological adaptations that could for instance allow them to out-compete bacteria in specific niches with low energy or nutrients availability [82]. However, previously studied localities within CCB displaying the same low energy stress, such as Pozas Rojas microbial mat, lack similar numbers of Archaea. Pozas Rojas exhibits extreme N:P ratios (156:1), and also fluctuating environments [33]. Despite all of the above, until AD were uncovered, archaeal abundances and diversity had not been reported before as a distinctiveness in any of the numerous explored sites within CCB [24, 27, 28, 35]. The abundance of Euryarchaeota members (halophiles and methanogens) may be related to the high-carbonate, salty-crust upper layer, and the black methanogenic bottom strata, acting as barriers on both sides on the mat.

The estimated α and β diversity indexes for the AD samples suggest resilience of the community, which is expected to be largely influenced by species diversity, as explained by the insurance hypothesis [83]. Our data shows a slight increase in archaeal diversity and richness indexes in dry seasons compared to the sample from the wet season (Table 2), possibly because that is the usual state of the AD site, since annual precipitations are not heavy enough to dissolve the salts and change the ecology of the mats (Supp. Table 1)

Another plausible explanation for an increased abundance and diversity in the Archaea domain in the AD mats is the *killing-the-winner* hypothesis [84], that proposes a negative frequency-dependent selection, in which bacterial types are affected by viral pressure, promoting the survival and viability of rare types, and thus maintaining high diversity. Viral reads in our AD metagenomes represent a relative abundance ranging from 23.3% to 31.6%, which is remarkably high compared to other metagenomes, and even higher than typical virus-rich environments, such as those obtained from samples of filtered sea water in South Korea [85], several deep sea sediments [86] and microbialites from SharkBay, Australia [71].

### 4.2 The rare biosphere in the AD community

Rare biosphere may represent a reservoir of genetic diversity that actively responds to environmental changes [87]. Given the predominance of an equitable rare biosphere in all the previously sampled sites of CCB [24], and the extreme and unpredictable conditions of the AD site, we considered it was important to explore rarity within the Archaea community.

By defining a “strictly rare” biosphere as the taxa with relative abundances below 0.01%, we found 50 archaeal OTUs. Within this group, 11 OTUs seem to be conditionally rare [67], maintaining their lower abundance in only one sampling time (the wet month), and reaching higher abundances in both dry samplings (Fig. 3A). Their dynamics lead us to speculate that those rare OTUs from the AD, along with the more abundant OTUs, could be benefiting of a saltier environment or drier conditions. In both cases, it is noticeable that although archaeal richness and overall abundance does not change much among samples, it is the rare (the one driven by environmental fluctuations) biosphere that exhibits variations. A similar response to cold versus warm conditions have been noticed in other systems within CCB [37].

Among this rare biosphere, some taxa were found to be transiently rare (i.e., absent in one or two of the samples) comprising 39 OTUs (Fig. 3B). We suggest that this type of rarity should be driven at least in part by stochastic processes, such as passive dispersal of lineages temporarily recruited from microbial seed bank, or due to immigration [88]. As an example, nine rare unclassified taxa that were not present or were in very low abundances in the initial sample AD1, became abundant in February sample (dry season) (Fig. 3; Suppl. Fig. 2). More metagenomic studies are in their way to explore such pattern in a longer time frame.

It should not be discarded that the lowest-abundant taxa of AD could be undergoing dormancy, a mechanism that maintains cells alive, but inactive and intermittently below detection thresholds [89]. Indeed, Archaea can enter a cellular dormant state [89–91], thereby providing an adaptive response to what would otherwise be a deleterious environmental perturbation. This has been experimentally studied analyzing the thermoacidophile archaeon *Metallosphaera prunae* that produces VapC toxins that drive cellular dormancy under uranium stress [92]. Another factor for dormancy in Archaea is predatory organisms, specifically, virus, which are abundant in the AD metagenomes (∼28%). As an example, a research group showed that rare and even inactive viruses, induce dormancy in the model archaeon *Sulfolobus islandicus* [90]. Also oligotrophy might be playing a role in Archaea dormancy, as dormant bacteria taxa have been found to be enriched in low phosphorus environment [89] and dormant microorganisms can also escape virus predation [93]. Considering the extremely skew stoichiometry, virus may be the drivers of the community structure, not only liberating immobilized nutrients to the system by lysing cells, but by maintaining a large panoply of rare taxa coexisting and avoiding predation.

### 4.3 Halophiles and methanogens Archaea in the AD

Halophiles and methanogens species were highly abundant in the AD, representing > 50% of the total Archaeal diversity. The relationship between halophiles and methanogens is well known, sharing a common ancestral habitat [78, 82, 94, 95] and both groups are found in microbial mats and microbial communities associate to precipitated minerals (endoevaporites) [70]. The methanogens living in the aforementioned environments need a high level of NaCl (0.5 M) for optimal growth [96] and usually are halotolerant or halophilic [97]. This type of hypersaline environments is very dynamic and wide spread in the world [98].

In our sampling site, halophilic and methanogenic Archaea are proposed to constitute the stable core of the AD during hypersaline conditions under the salt crust. Accordingly, other studies indicate that salinity is the major abiotic factor that allows the shaping of microbial communities, especially in sediment surfaces, stromatolites, hydrothermal vents, hypersaline mats and anoxic saline water. These studies have demonstrated that saline sediments contain communities with higher unique biodiversity values in comparison with other environments [99]. Other studies demonstrate that changes in salinity, sulfate and availability of substrates could possibly stimulate the production of methane [70].

The most abundant OTUs in the analyzed AD metagenomes were similar to *Methanohalophilus mahii*, a halophilic-methanogen Archaea described as a non-marine methanogen, adapted to hypersaline environments; its metabolism requires 1.0-2.5 M NaCl for optimal growth and methanogenesis, using methanol and methylamines as substrates [95]. Other OTUs from AD were related to *Natrialba taiwanensis, Haloferax sulforifontis, Methanoplanus petrolearius* and *Haloterrigena thermotolerans*. Halophilic and methanogen species coexisted within the microbial mat, along with some unexpected genera belonging to Euryarchaeota phylum. Among the latter we can find, *Thermococcus, Natronomonas, Picrophylus, Archaeoglobus, Aciduliprofundum* and even *Thermoplasma,* a lineage that lacks a cell wall. These microorganisms are mostly reported from hydrothermal vents, displaying a high level of tolerance to low and high temperatures and a wide range of pH [100, 101]. The presence of these taxa along with the previous observations, adds evidence to our proposal stating that CCB microbial communities have both marine and magmatic affinities [30, 32]. Therefore, the Archaea that we observed in AD are witness of both environments, the deep aquifer propelled by magmatic heat within the mountain of San Marcos y Pinos [29], and the surface life, where photosynthesis occurs, rarely, after rainy days. Hence, part of the AD diversity might have emerged from the deep sediments that contained the mineral conditions of the ancient ocean [52].

### 4.4 Thermophilic and other related Archaea lineages in the AD

Summing to the hypothesis of magmatic influence to the microbial community of this unusual site, Crenarchaeota taxa are consistently present in the AD mat. These include thermophilic genera such as *Thermoproteus, Caldivirga, Ignicoccus, Sulfolobus, Pyrolobus,* among others. In addition, Thaumarchaeota members were detected, such as “*Candidatus* Nitrososphaera gargensis”, *Nitrosoarchaeum limnia*, and Marine Group 1 Thaumarchaeota. The presence of two OTUs were unexpected: *Cenarchaeum symbiosum,* a psycrophillic archaeon, previously reported as a symbiont of a several sponge species [102], and *Nitrosopumilus maritimus,* considered ubiquitous on oligotrophic oceans [103], another AD finding consistent with the marine signatures frequently found in CCB microbes [25, 28, 30, 44, 53]. The same occurs with *“Candidatus* Korarchaeota cryptophilum”, a representative of Korarchaeota phylum, its presence in AD is related to the marine origin of CCB and an incoming colonization of geothermal-terrestrial environments, a feature shared among the thermophiles [104].

Both Crenarchaeota and Thaumarchaeota have in general terms similar metabolic capabilities. PCR-amplified *amoA* genes from DNA of each sample (AD1, AD2, AD3; data not shown) provided evidence of ammonia oxidation capabilities as potential energy source and nitrification [105], suggesting a role for chemolithotrophy of these taxa in CCB, using a large panoply of organic compounds as well as CO_2_, iron, nitrogen and sulfur compounds as electron acceptors, as for instance methanobacteria (Euryarchaeota) and hyperthermophilic Crenarchaeota members use sulfur as an electron acceptor [106].

Other unexpected but interesting member of the phyla in the AD metagenomes were the Nanoarchaeota *Nanoarchaeum equitans,* a small-sized archaeon and an ectoparasitic relationship with the Crenarchaeota *Ignicoccus*, which was also detected in the sample [107]. Nanoarchaeaota phylum has a widespread habitat distribution with diverse physicochemical features compatible with hot springs and other mesophilic hypersaline environments [108].

## 5. Perspectives and conclusions

The Archaean Domes microbial mats at the oasis Cuatro Ciénegas Basin, were recently discovered as a hypersaline extreme site. In these mats we uncovered through metagenomic analysis, one of the highest diversities registered so far in the Archaea domain, considering the site’s geographic small scale. Most of the 230 OTUs observed in this unusual small shallow temporal pond, are part of the rare biosphere and form a stable core community. Within this core, we observed halophiles and methanogens, but also spatially unexpected archaeal taxa, that thrive under high salt concentrations. We also observed a transient rare biosphere that appears to be enriched under dry environmental conditions, suggesting seasonal dynamics shaping community assemblage. In order to explore this group of taxonomic the unclassified rare taxa more carefully, we are in the process of analyzing more metagenomes in different seasons as well as manipulating the environment using mesocosms experiments. More sequencing effort in deep sediment cores will also help to look for the deep anaerobic biosphere, as well as eliminate blind spots in phylogeny of unclassified Archaea, and this will require differential coverage binning approach [109] using all available metagenomes from the AD to describe phylogenetic novelty within AD at CCB.

This highly diverse ecosystem within Cuatro Cienegas, Mexico, arises as an attractive novel site for evolutionary, ecological, astrobiological and bioprospecting studies. The AD is, so far, the most diverse microbial community found in CCB, despite its extreme conditions. Since this area is being subjected to intense water exploitation by agricultural practices, and desiccation has become a common occurrence in numerous ponds, it is priority for our research group to keep investigating the ecology of the adaptation of this highly diverse archaeal-rich microbial communities to fluctuating temperature and rainfall conditions, while working with shareholders on changes in policy of water usage.

Now proven as an archaeal rich extreme site, CCB is once more attracting attention as an astrobiological model [38, 41]. The Archaean Domes not only can take us further back into the “lost world” but it is also a site that keeps providing evidences and new keys to understand how “life cycle” could have been originated on Earth, or (will be) possible on, for example, Mars.

## Conflicts of interest

The authors declare that there is no conflict of interest regarding the publication of this paper.

## Acknowledgments

We want to thank to Hamlet Aviles-Arnaut, Gabriel Moreno-Hagelsieb for their critical review of the manuscript. Also, thanks to Hamlet Avilés-Arnaut, Irene Ruvalcaba-Ortega and Ricardo Canales-Del Castillo for their valuable technical support and critical observations throughout the project. Thanks to Kendra Rivera and Josué Corona for their technical help on the experiments. Finally, we thank SEMARNAT and APFF Cuatro Ciénegas for facilitating the sampling and in particular Rancho Pozas Azules, PRONATURA Noreste for access and permission to sample in the CCB Natural Protected Area.

## Funding Statement

We thank Universidad Autónoma de Nuevo León for funding field work through the PAICYT program granted to Susana De la Torre-Zavala during 2015. We thank the Alianza WWF-Fundación Carlos Slim fund to Valeria Souza and Luis E. Eguiarte.

## SUPPLEMENTARY MATERIAL

**Supp. Table 1.** Physicochemical parameters measured for Archaean Domes on April 2016. IBV (volts); Temp: temperature °C; SPC: specific conductance (μS/cm); Salinity (psu); TDS: total dissolved solids (g/L); pH (0-14); LDO: dissolved oxygen (mg/L); CH4 (μg/L).

| Physicochemical characteristics of AD | IBV | Temp | SPC | Salinity | TDS | pH | LDO | CH<sub>4</sub> | CO<sub>2</sub> |
| --- | --- | --- | --- | --- | --- | --- | --- | --- | --- |
| Water sample | 7.7 | 31.45 | 77.21 | 53.36 | 49.62 | 9.94 | 78.4 | 12.4 | Not detected |
| Mat Sample | 8.0 | 34.35 | 77.31 | 52.55 | 48.76 | 9.75 | 1: 165.3<br>2: 7.3 | N/A | Not detected |

**Supp. Fig. 1.**
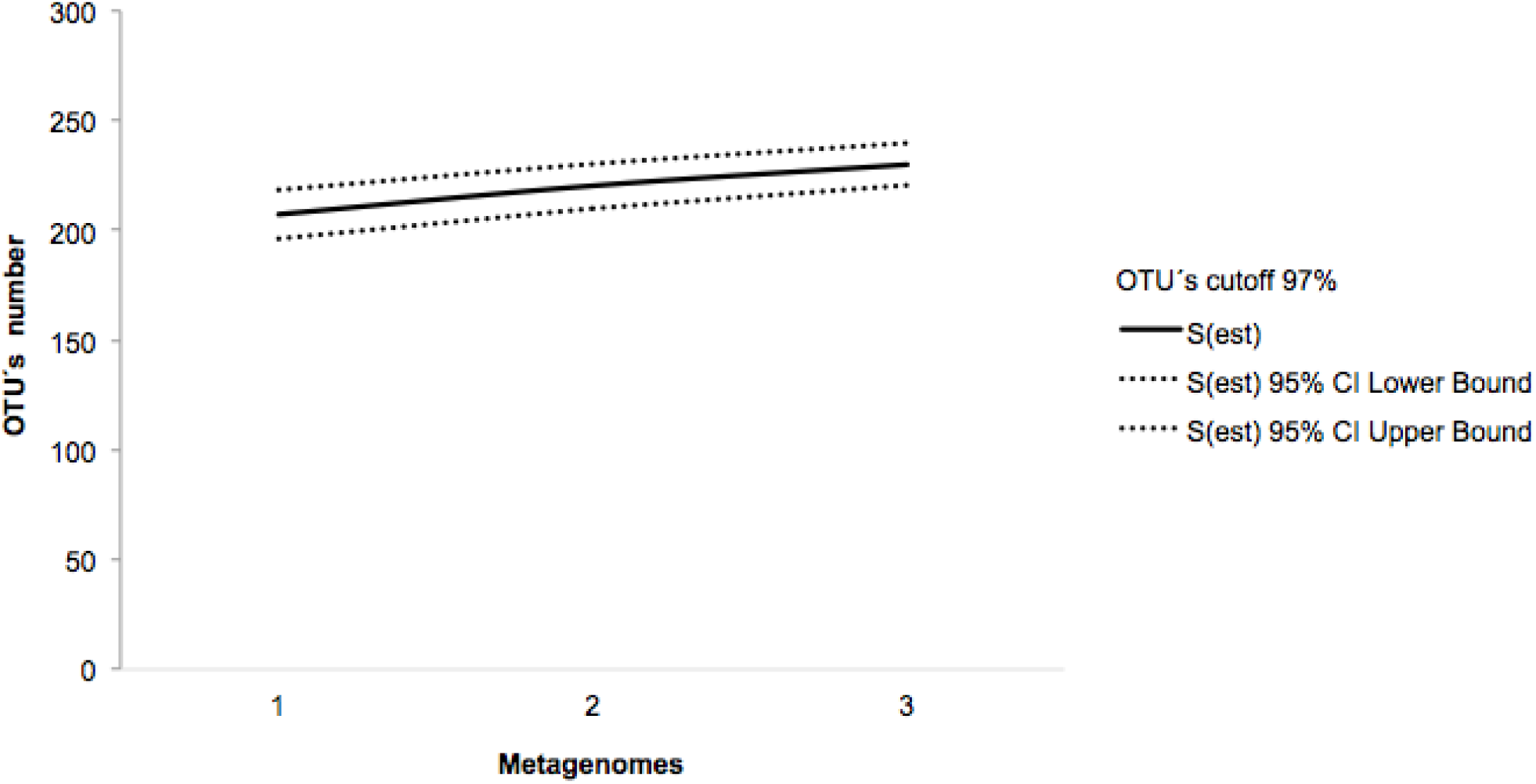
Rarefaction curve to estimate the richness of Archean Domain in the different season samples.

**Supp. Fig. 2.**
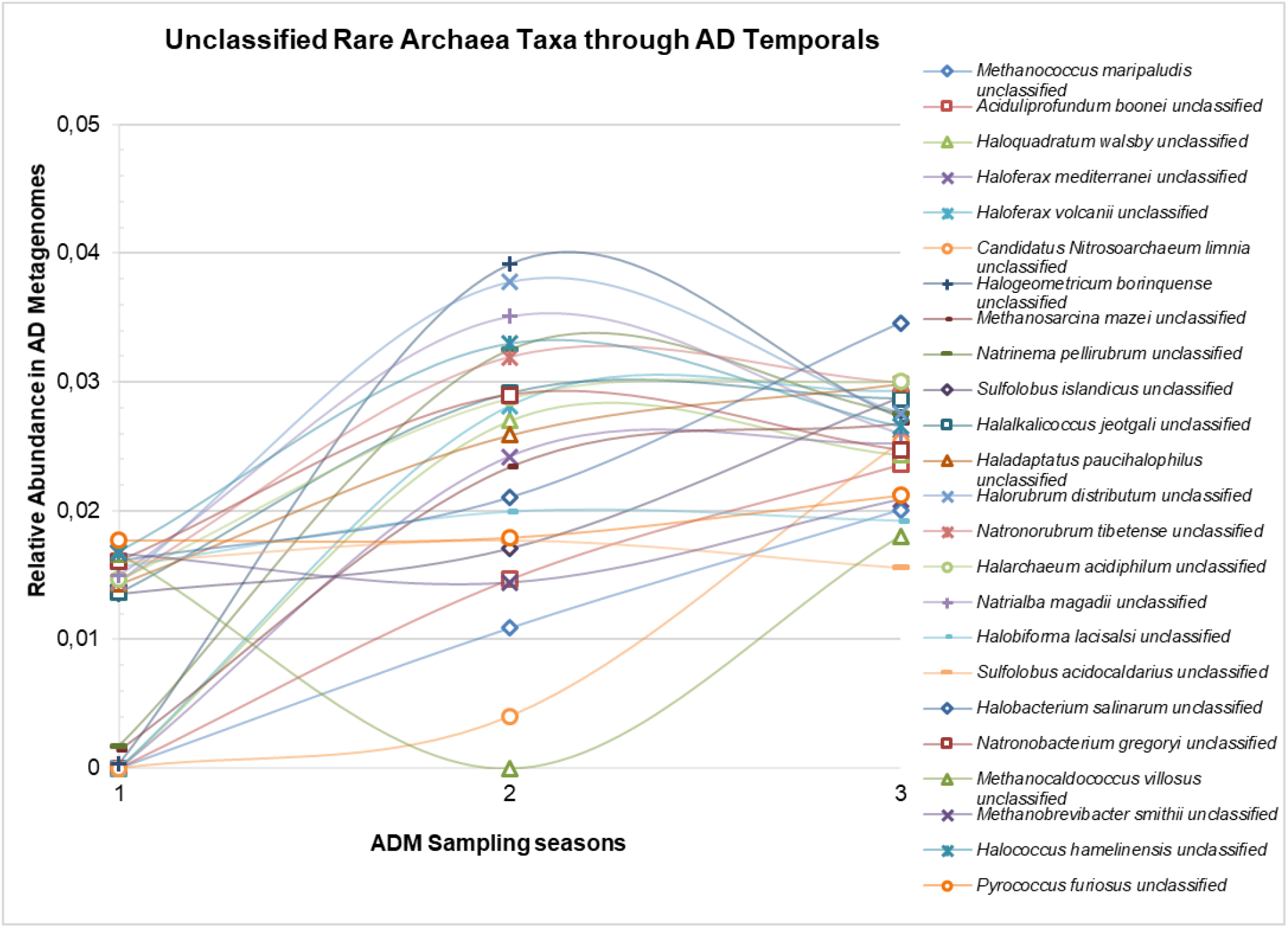
Unclassified Rare Archaea Taxa in the AD.

## Notes

https://www.mg-rast.org/linkin.cgi?project=mgp90438

